# Addendum to Ancient DNA data from Mengzi Ren, a Late Pleistocene individual from Southeast Asia, cannot be reliably used in population genetic analysis

**DOI:** 10.1101/2025.03.24.645126

**Authors:** Daniel Tabin, Nick Patterson, Matthew Mah, David Reich

**Affiliations:** Department of Human Evolutionary Biology, Harvard University, Cambridge, Massachusetts 02138, USA; Broad Institute of MIT and Harvard, Cambridge, Massachusetts 02142, USA; Department of Genetics, Harvard Medical School, Boston, Massachusetts 02115, USA; Howard Hughes Medical Institute, Boston, Massachusetts 02115, USA

## Abstract

In addition to the issues pointed out in Tabin et al^1^, the MZR data from Zhang et. al 2022^2^ are suggestive of high levels of contamination from a source similar to modern Han Chinese, the majority population in the country where MZR was sequenced. In fact, MZR can be modeled entirely as Han-related ancestry and noise. These results raise further concerns about the veracity of the MZR data and thus the paper’s historical conclusions.

---

One of Zhang et al.’s most surprising claims is that Northeast Asians including Han Chinese share more alleles with the DNA sequences obtained from the remains of the MZR individual sampled from Late Pleistocene Southeast Asia than they do with later ancient individuals from early Holocene Southeast Asia^2^. We replicate this finding with symmetry statistics^3^ of the form *f*_*4*_*(MZR, Early Holocene Southeast Asian*^*4-8*^; *Han*^*9,10*^, *Mbuti*^*9,10*^) (Supplementary Data 1), of which many are significantly positive (and none are significantly negative), using a dataset of 8.65 million single nucleotide polymorphisms (SNPs) which we ascertained to increase statistical power^1^. However, we were concerned that the signal of MZR affinity to Northeast Asians might not reflect the true ancestral heritage of MZR, and instead might be indicative of contamination. That would be especially problematic, because the contamination would also produce a false affinity towards Native Americans. This could falsely generate some of the findings of Zhang et al^2^.

## MZR’s nuclear DNA are consistent with being largely Han Chinese contamination

We were concerned that the signal of MZR affinity to Northeast Asians might not reflect the true ancestral heritage of MZR, and instead might be an artifact of contamination from modern people with Northeast Asian ancestry like Han, the majority population in China where MZR was sampled and sequenced.

The data in fact point to very high rates of Han Chinese-related contamination. When we measure shared drift of MZR and other ancient Southeast Asians with diverse modern Eurasians using the statistic f_3_(Ancient Southeast Asian; Modern Eurasian, Yoruba), MZR shares the most drift with Han and populations closely related to Han (Supplementary Data 6). While the ∼8000 year old Ancient Southeast Asian individual from Liangdao also shares an affinity to Han, it lacks the affinity to Northeast Asians such as Korean and Daur that MZR (and Han) have. The ∼12000 year old Longlin, the ∼8000 year old Hoabinhian, the ∼7000 year old Sulawesi hunter-gatherer, and the ∼40000 year old Tianyuan individual all share less drift with Han^4,5,7^. The Longlin, Hoabinhian and Sulawesi individuals also all share more drift with Australasian populations such as Onge, Papuans, and Australians.

To understand the genetic affinities of MZR’s genome, we used the software *qpAdm*^*11*^, attempting to model MZR as a mixture of Han and French (representing plausible contaminants), and a dummy population consisting of heterozygous genotypes at each SNP to represent the high error rate we have shown exists in MZR^1^ (we hypothesize that sequencing error might randomly switch each genotype at each SNP to the alternate allele, hence the idea to model error as a 50-50% frequency at each SNP). *qpAdm* requires using a set of “Right” reference populations that are differentially related to the different ancestry sources, making it possible to tease them apart, and for this purpose we used Yoruba, Papuans, Onge, Mala, Karitiana, Japanese, Georgian, and Ami. Due to the fact that MZR had most of its unusual errors on the ends of its sequences (Figure 1), we carried out analyses with three different trimming schemes. First, we analyze the full sequences (untrimmed). Second, we analyzed sequence trimmed 8 bases on both ends.

Finally, we trimmed 2 bases off the 5’ end and 17 bases off the 3’ end of sequences, following the procedure of Zhang et al^2^. All three trimming frameworks provided qualitatively identical results with quantitatively similar numbers.

Focusing here on the results based on applying *qpAdm* to the sequences with the last 8 bases trimmed, we find that MZR can be fit well as having 90.1±0.9% Han Chinese- related DNA with the remaining ∼9.9% coming from the dummy population capturing the unusually high sequencing error in the data. This model fits even when moving the French to the outgroup populations indicating no evidence of European contamination (however, both the Han and noise sources are required). This type of model does not fit for other ancient samples including a Taiwanese individual from Liangdao (using the individual with lower coverage that MZR, thus showing that the fit of MZR is not an artifact of low coverage which increases the chance of a model passing), Longlin, the hunter gatherer from Sulawesi, the Hoabinhian from Laos, or Tianyuan. A non-fitting model is what is expected if an ancient individual has ancestry from a lineage that is phylogenetically more closely related to Papuans, Onge, Ami, or any other Right population than to either Han or French (Table 1). In contrast, a fit is consistent with all data being Han-related contamination plus error.

**Table 1:**
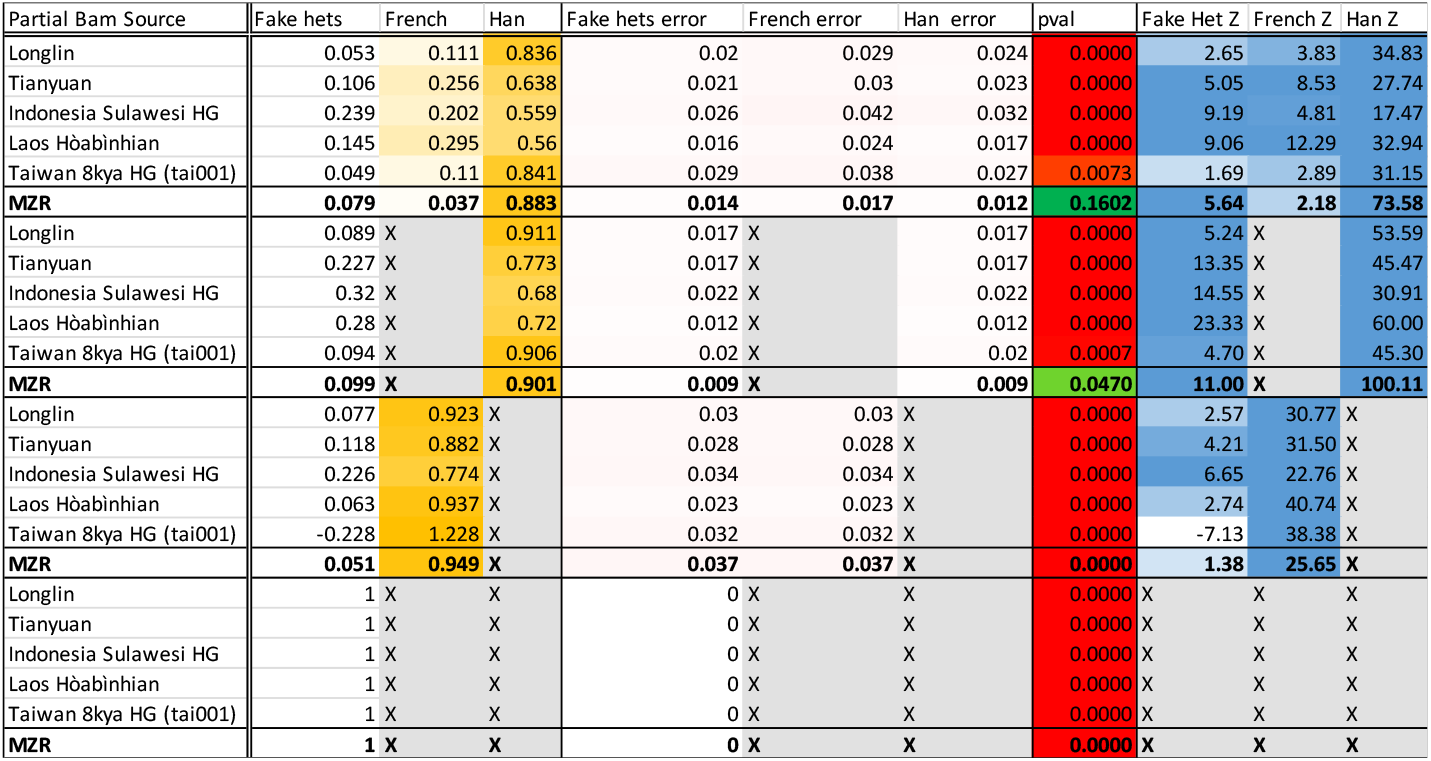
*qpAdm* results when modeling ancient Southeast Asians as mixtures of Han Chinese, French, and a dummy population meant to simulate the effects of sequencing error. When modeling ancient Southeast Asian samples as mixtures of Han, French, and a dummy population used to represent sequencing error, with many plausible modern representatives of Southeast Asian ancestry in the right population set, all targets other than MZR fail to be properly modeled. Unlike other published ancient Southeast Asians, MZR is modeled as 90.1% derived from modern Han Chinese and 9.9% error and fails to be properly modeled when Han Chinese is removed from the sources. This is an effect specific to Han, as when French is removed, MZR continues to produce a fit. These results are consistent with MZR’s data being almost entirely Han contamination, and unlike all the other ancient individuals, provides no evidence of genuine ancient Southeast Asian ancestry.

## MZR’s high error rates makes separating out genuine sequences nearly impossible

In order to extract potentially genuine MZR sequences, we divided the MZR data based on the rate of DNA damage on the sequences. We used *PMDtools*^12^ to infer if sequences are not likely to be damaged and thus likely to be authentic. Given the three groups of PMD values sequences tend to fall into (Figure 2), we split the sequences into three parts. The sequences in the “negative” group with PMD scores < 0 have the lowest likelihood of being authentic ancient DNA. The “low” group includes sequences with PMD scores between 0 and 2.7 consisting of sequences that have characteristic ancient DNA damage but not in the most diagnostic positions. The “high” group sequences have the highest PMD scores >2.7 with the strongest evidence of characteristic ancient DNA damage. We computed f_4_-statistics such as *f*_4_(*MZR div 1, MZR div 2;Han,French*) using the *qp4diff* program in ADMIXTOOLS^3^ as the difference between two statistics:

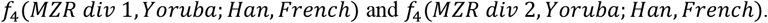

**Figure 2:**
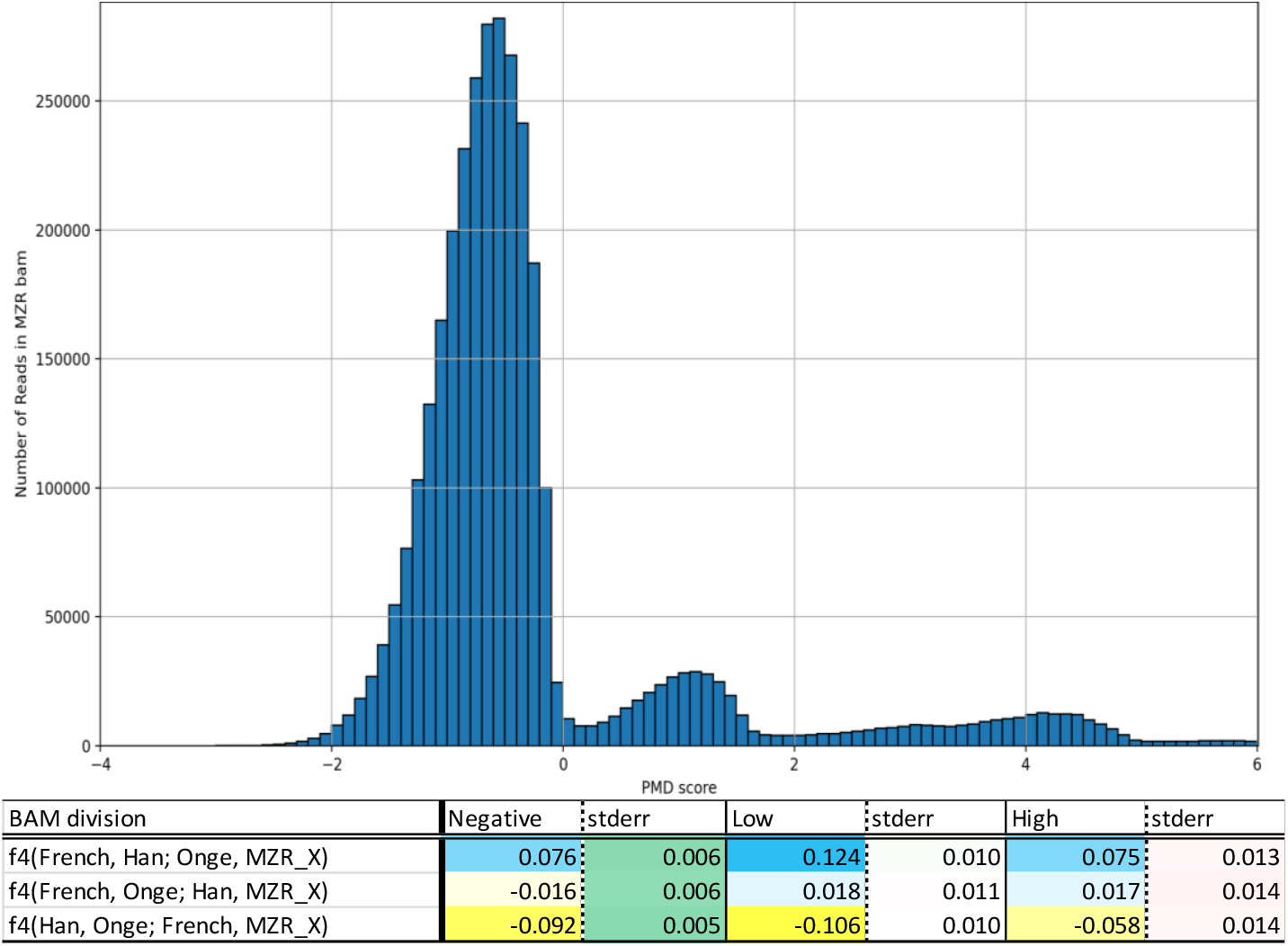
MZR subdivisions show differential relatedness toward French, Onge, and Han. (a) A histogram of sequences from the MZR bam by PMD-score per sequences. MZR bams are binned into 0.1 PMD score difference bins. Three modes are observed: one with negative damage, one with low but positive damage, and one with high damage. (b) Computing the three unique f_4_ statistics consisting of Han, Onge, French, and an MZR subdivision produces different population genetics results depending on the subdivision used.

Because *qp4diff* does not require restriction to positions covered by sequences in two subdivisions, which is only a tiny fraction of sites for the low coverage MZR data, we can use much more of the data and obtain a more precise estimate of the statistic. We find that sequences with moderate evidence of damage have significant excess attraction to Han when compared both to the sequences with the strongest evidence of damage (Z > 6.89), and the sequences with the least evidence of damage (Z > 9.23). A full comparison of the MZR divisions to Han and other Eurasians can be found in Supplementary Data 6.

These results are not consistent with analyzable population genetic data, both due to MZR’s relative affinities to Han and French differing based on damage-score, and the fact that this differential affinity is not typical of contaminated data. While the sequences with the least evidence of damage are expected to show affinity to Han if MZR is contaminated, it is surprising that the least damaged sequences also show affinity to Han.

A potential explanation for this is the strange mismatch patterns on the ends of MZR sequencess^1^. Whatever processes generated the errors—which appear across mutation classes and occur more on the 3’ end than 5’ end—also affected C->T and G->A mismatches, which are used to identify genuine ancient DNA through damage restriction. This means that it is not possible to separate the sequences into contaminated and uncontaminated bins reliably, making it difficult to extract “genuine” MZR sequences from the presumably contaminated data.

## Discussion

The MZR data have specific affinity to modern Han Chinese and other related populations. This is *a priori* surprising given the date and location of the MZR individual. This affinity is so strong that MZR can be well modeled as entirely Han with added noise. Neither of these traits are shared with other ancient Southeast Asians and both raise additional concerns regarding the reliability of the MZR data. Contamination seems more plausible than a population with Han-like ancestry existing in Yunnan province 14 thousand years ago.

## List of Supplementary materials

**Supplementary Data 1:** f-statistics of the form f_4_(MZR, Ancient Southeast Asia; Han, Outgroup)

**Supplementary Data 2:** Table of f_3_-outgroup showing shared drift between ancient Southeast Asians and modern Eurasians

**Supplementary Data 3:** Table of f_4_-differences comparing MZR subdivisions to Han and other modern Eurasians

## Acknowledgements

We thank Xiaoming Zhang and co-authors for collegial discussions, which informed the final manuscript. We thank three anonymous reviewers of earlier versions of this manuscript. This research was supported by NIH grant (HG012287), the John Templeton Foundation (grant 61220), and by the Howard Hughes Medical Institute.

